# The histone H3.3 K27M mutation found in diffuse midline gliomas coordinately disrupts adjacent H3.3 Ser31 phosphorylation and the fidelity of chromosome segregation

**DOI:** 10.1101/2022.05.27.493485

**Authors:** Charles A. Day, Florina Grigore, Faruck L. Hakkim, Alyssa Langfald, Sela Fadness, Paiton Schwab, Leslie Sepaniac, Jason Stumpff, David J. Daniels, Kevin T. Vaughan, James P. Robinson, Edward H. Hinchcliffe

## Abstract

During the cell cycle, differential phosphorylation of select histone H3 serine/threonine residues regulates chromatin structure, necessary for both dynamic transcriptional control and proper chromosome segregation^1-2^. Histone H3.3 contains a highly conserved serine residue (Ser31) within its N-terminal tail that is unique to this variant. During interphase phosphorylation of Ser31 amplifies stimulation-induced transcription and is required for early metazoan development^3-6^. During mitosis Ser31 phosphorylation at the pericentromere supports proper chromosome segregation, albeit by unknown mechanisms^7-10^. H3.3 Ser31 is flanked by mutational sites that drive several human cancers, including pediatric gliomas^5-8^. This is typified by the H3.3^K27M^ mutation found in ∼80% of diffuse midline gliomas, which undergo epigenetic reprogramming in proliferative cells coordinate with loss of global H3 lysine 27 trimethylation (H3K27Me3)^11-14^. However, whether the K27M mutation influences the neighboring Ser31 phosphorylation and whether disrupting Ser31 phosphorylation plays a distinct role in driving gliomagenesis has not been tested. Here we show that H3.3^K27M^ mutant cells have reduced capacity for H3.3 Ser31 phosphorylation at the mitotic pericentromere, increased rates of chromosome missegregation, and impaired G_1_ checkpoint responses to chromosome instability. CRISPR-reversion of K27M to wild-type restores phospho-Ser31 levels and suppresses chromosome segregation defects. CRISPR editing to introduce a non-phosphorylatable H3.3^S31A^ alone is sufficient to increase the frequency of chromosome missegregations. Finally, expression of H3.3^S31A^ in a PDGFβ-driven RCAS/TVA mouse model is sufficient to drive high grade gliomagenesis, generating diffuse tumors morphologically indistinguishable from those generated by H3.3^K27M^ expression. Importantly, this occurs without the loss of H3K27 triple methylation that is the hallmark of K27M tumors. Our results reveal that the H3.3 K27M mutation alters the neighboring Ser31 phosphorylation, and loss of proper H3.3 Ser31 phosphorylation contributes to the formation of diffuse midline gliomas.

---

Diffuse midline gliomas with the H3^K27M^ mutation, including diffuse intrinsic pontine gliomas (DIPG), are amongst the most aggressive primary malignant brain tumors in children^10-11^. Despite extensive research and >100 clinical trials, median survival for these patients after diagnosis and radiation therapy remains ∼1 year^15-16^. Pioneering genetic analysis of diffuse midline gliomas identified heterozygous somatic missense mutations in the H3F3A gene that encodes the histone variant H3.3, the most prominent mutation being lysine 27-to-methionine (K27M)^11-14^. This mutation acts as a dominant negative and is sufficient to drive dramatic decreases in H3 Lys27 di- and tri-methylation (H3K27Me2 and H3K27Me3) on all histone H3 proteins, both mutant and wild-type throughout the genome^13-14^. The loss of these transcriptionally repressive marks during periods of cell proliferation results in global epigenetic reprogramming and is thought to maintain dividing DIPG cells in a stem-like state^17-19^. While K27M mutations in canonical H3.1 genes (lacking Ser31) are also found in DIPGs, these are far less common, the tumors less aggressive, and those tumors often exhibit an early secondary mutation resulting in the loss of *CDKN2A* expression and disrupted cell cycle checkpoint control^11-12^.

In addition to epigenetic reprogramming, diffuse midline H3.3K27M mutant tumors also exhibit Copy Number Alterations (CNAs), suggesting these tumors are chromosomally unstable^21-22^. Chromosome instability is driven by complete chromosome missegregation to one daughter cell or by chromosomes that do not completely disengage and separate during anaphase – causing bridging and breakage of large chromosome regions (such as arms). These arms can be lost to the cell, maintained as extra chromosomal DNA (found in many gliomas), or become translocated to another chromosome by break and repair pathways^22-24^. Importantly, chromosome missegregation during mitosis is sufficient to drive the extensive structural chromosomal variations that resemble the genomic features commonly associated with tumorigenesis ^24^.

Chromosome missegregation is normally monitored by checkpoint mechanisms that center on both the centromere/kinetochore and pericentromere regions of mitotic chromosomes^25^. During mitosis histone H3.3 is phosphorylated at residue Ser31, which is one of five amino acid changes that differentiate H3.3 from canonical H3.1^26^. Importantly, mitotic histone H3.3 Ser31 phosphorylation in mammals begins during prometaphase/metaphase^7,10^, where it is restricted to the pericentromeric heterochromatin (PCH); Ser31 then becomes dephosphorylated as the disjoined sister chromatids regress during anaphase (Figure 1A). However, chromosomes which fail to properly congress during metaphase or lag in the spindle midzone during anaphase have extensive Ser31 phosphorylation that spreads along their chromosome arms (Figure 1B).

**Figure 1.**
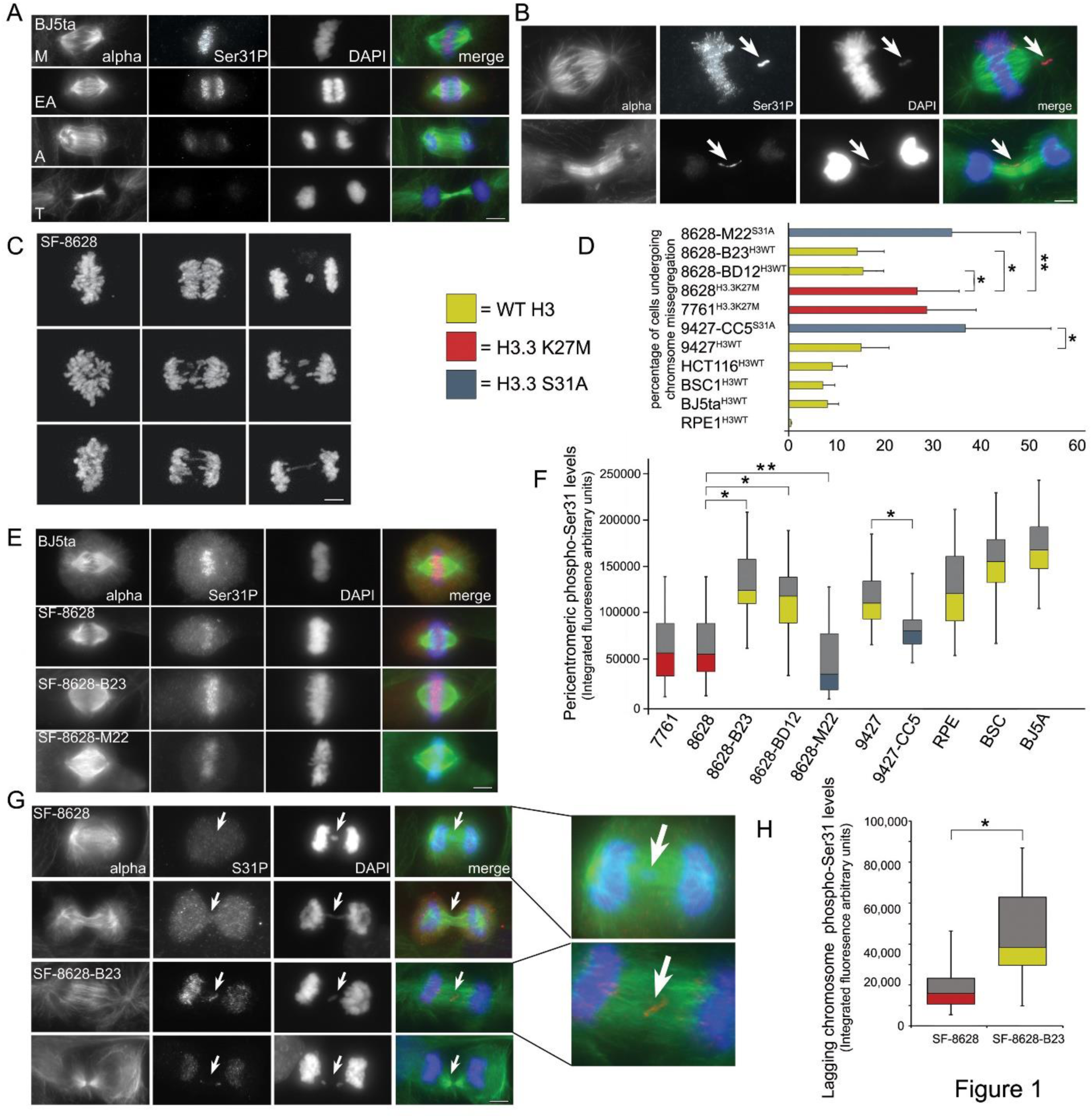
H3.3K27M mutation alters Ser31 phosphorylation and chromosome segregation. **A**. Distribution of H3.3 Ser31 phosphorylation during metaphase-anaphase transition in normal human cells. Phospho-Ser31 is restricted to the pericentromere region in metaphase (M) and early anaphase (EA) cells, then rapidly becomes dephosphorylated as anaphase (A) progresses into telophase (T), and the nuclear envelop reforms. **B**. Lagging chromosomes before anaphase (upper panels) or after anaphase (lower panels) have high levels of phospho-Ser31 along their arms (arrows). **C**. Frames from live-cell microscopy sequences showing chromosome missegregation patterns in patient-derived SF-8628^H3.3K27M^ pediatric diffuse midline glioma cells expressing H2B-GFP and undergoing anaphase. **D**. Percentage of anaphase cells imaged by live-cell spinning disk confocal microscopy with one or more lagging or bridged chromosome, color-coded for histone H3 genotype: H3^WT^ (yellow: RPE-1, BJ5ta, BSC-1, HCT-116, SF-9427, SF-8628-B23, and SF-8628-BD12), H3.3^K27M^ (Red: SF-7761 and SF-8628) or H3.3^S31A^ (Blue: SF-9427-CC5 and SF-8628-M22). *n* = 50 cells for each. The reversion of M27 (mutant) to K27 (WT) results in a significant decrease in lagging chromosomes, while the insertion of S31A along with M27K reversion does not. Graph shows mean from five separate experiments ± SD. \**P* < 0.05, \*\**P* ≥ 0.05 two-tailed *t*-tests. **E**. Pericentromeric H3.3 Ser31 phosphorylation in metaphase cells with H3^WT^ (BJ5ta and SF-8628-B23), H3.3^K27M^ (SF-8628) or H3.3^M27K/S31A^ (SF-8628-M22). **F**. Box and Whisker plots (median value, 50^th^ and 75^th^ percentile, Max and Min) of pericentromeric H3.3 phospho-Ser31 levels measured from SF-7761, SF-8628, SF-8628-B23, SF-8628-M22, SF-9427, SF-9427-CC5, RPE-1, BSC-1, and BJ5ta cells. *n* = 40 cells for each from four separate experiments. \**P* < 0.05, \*\**P* ≥ 0.05 two-tailed *t*-tests. **G**. H3.3^K27M^ results in decreased phospho-Ser31 on lagging chromosomes (arrows) in post-anaphase cells. **H**. Box and Whisker plots (median value, 50^th^ and 75^th^ percentile, Max and Min) of H3.3 phospho-Ser31 levels on lagging chromosomes measured from SF-8628 vs SF-8628-B23. *n* = 30 cells for each. \**P* < 0.05 two-tailed *t*-test.

Ser31 neighbors Lys27, and this amino acid substitution changes the topology of the histone H3.3 tail, which may impact Ser31 phosphorylation levels and the regulation of chromosome segregation. To test whether the K27M mutation affects chromosome segregation patterns, we directly measured the frequency of chromosome missegregation events using live-cell imaging of the metaphase-anaphase transition in a panel of cells expressing GFP-histones as fluorescent chromosome markers. The presence of one or more lagging and/or bridged chromosomes is easily detected using this system, and these are scored as missegregated (Figure 1C). We examined cell lines with wild-type histone H3, including the normal diploid cell lines RPE1, BJ5ta, and BSC1, a chromosomally stable human colon tumor cell line (HCT-116), and the human pediatric glioblastoma cell line SF-9427. These cell lines all exhibited low levels of chromosome missegregation (<10% of mitotic divisions resulted in a missegregated chromosome), with the exception of the pediatric GBM cell line SF-9427, which shows a modest increase (∼15%) in chromosome instability (Figure 1D). However, when we analyzed two patient-derived K27M-mutant pediatric diffuse midline glioma cell lines (SF-8628 and SF-7761) we observed chromosome missegregation events occurring at significantly higher rates, with missegregations in the SF-8628 cells occurring in ∼27% and in ∼29% of SF-7761 cells analyzed (Figure 1C, D).

To test whether these chromosome segregation defects are in direct response to the K27M mutation, we used CRISPR (clustered regularly interspaced short palindromic repeats) gene editing of SF-8628 cells to revert them to wild-type for histone H3.3. We isolated two independent clones: SF-8628-B23 or ”B23 cells”, where the mutant HF3F3A allele becomes deleted (HF3F3A^WT/null^), and SF-8628-BD-12 or ”BD-12” cells, which have a M27K insertion in the mutant allele rendering these cells HF3F3A^wt/wt^. To generate these indels, cells were transfected with guide RNAs to facilitate the reversion, along with soluble Cas9 protein. This allows CRISPR-editing to occur, but the Cas9 protein is rapidly degraded, ensuring that by the time of clonal selection there is no continued CRISPR editing that could cause off-target effects or chromothripsis. After CRISPR reversion B23 and BD-12 cells are viable and grow to confluency; they lose their anti-K27M immuno-reactivity and regain H3 Lys27 trimethylation levels comparable to WT cells (Supplemental Figure 1). Live-cell analysis of mitotic division reveals a significant decrease in chromosome missegregation from ∼27% of the parental SF-8628^K27M^ anaphase cells to ∼14% for the B23^M27K^ and ∼15% for the BD-12^M27K^ anaphase cells respectively (Figure 1B, D). To determine whether these chromosome missegregation defects can be affected by downregulating Ser31 phosphorylation directly, we used CRIPSR-editing of SF-8628 cells to create a double mutant allele that reverts the K27M mutation to WT by insertion, while also introducing a non-phosphorylatable S31A mutation. These cells (SF-8628-M22 or “M22 cells”) also grow to confluency, indicating they are viable (Supplemental Figure 1). Importantly, M22 cells exhibit increased chromosome missegregation at levels (∼33%) comparable to the parental mutant SF-8628 cells (Figure 1D). We also used CRISPR editing to introduce the S31A mutation into the H3^WT^ SF-9427 pediatric GBM cell line (SF-9427-CC5 cells or “CC5 cells”). Chromosome missegregation is increased in these H3.3^S31A^ expressing CC5 cells as well (37%), compared to the parental SF-9427 cell line (Figure 1D). Together these data suggest that proper regulation of phospho-Ser31 levels is required to maintain accurate chromosome segregation during anaphase, and that the presence of the K27M mutation results in an increase in chromosome missegregation events in DIPG cells.

To test whether the K27M mutation directly influences mitotic Ser31 phosphorylation levels, we measured Ser31 phosphorylation in individual cells by quantitative fluorescence microscopy. Direct measurement of mitotic cells revealed that the presence of the heterozygous H3.3 K27M mutation decreased median mitotic Ser31 phosphorylation by ∼2-3 fold compared to H3 wild-type cells (Figure 1E, F). We analyzed our CRISPR–edited H3.3 wildtype revertant cell lines (SF-8628-B23^H3.3M27K^ and SF-8628-BD-12^H3.3M27K^) and found that levels of Ser31 phosphorylation increase in the absence of the K27M mutation. The M27K revertant cells have mitotic Ser31 phosphorylation levels restored to those comparable to WT human cells and H3.3 WT glioblastoma cells (Figure 1F). When we examined our CRISPR-edited S31A cell lines (SF-8628-M22 and SF-9427-CC5) we found that the introduction of an S31A non-phosphorylatable mutant allele resulted in significant loss of overall mitotic Ser31 phosphorylation in metaphase cells compared to their respective parental cell lines.

Previously we have shown that Ser31 becomes hyperphosphorylated in response to chromosome missegregation, and phosphorylation spreads to the arms of the missegregating chromosome, and ultimately to both daughter nuclei^10^. This is correlated with the stabilization of nuclear p53 in these daughter nuclei and cell cycle arrest in both daughter cells, as previously shown^10, 26-27^. We also found that the accumulation of nuclear p53 in response to chromosome missegregation is prevented by masking phospho-Ser31 by antibody microinjection^10^. Together, our previous work suggested a model in which chromosome mis-positioning during anaphase triggers H3.3 Ser31 hyperphosphorylation on the lagging chromosome which spreads to both daughter nuclei. This triggers the induction of nuclear p53 in response to chromosome missegregation and serves to pause proliferation in both of these newly-generated aneuploid daughter cells^10, 27^. Here we examined whether the K27M mutation can also alter Ser31 phosphorylation on lagging chromosomes, using our matched pair of SF-8628 cells. Consistent with our measurements of metaphase Ser31 phosphorylation levels, we find a significant decrease in phospho-Ser31 on chromosomes lagging in the spindle midzone in the K27M-expressing cells (Figure 1G, H). The loss of Ser31 phosphorylation is reversed when the K27M mutation is lost in the B23 cell line. This suggests that K27M cells may have defective checkpoint signaling to respond to chromosome missegregation.

DIPG cells are transformed and have a high mutational load^11, 12, 20, 21^. Even in our revertant cell lines, the SF-8628-B23 and BD-12 cells continue to missegregate chromosomes at a higher frequency than WT cells. Therefore, we tested whether H3.3 mutations alone, expressed in normal, diploid cell lines can induce chromosome segregation errors. To do so we transiently expressed either K27M-GFP or S31A-GFP in BJ5ta and BSC1 normal diploid cells. Three days after introduction of the transgene we analyzed individual cells progressing into anaphase by live imaging (Figure 2A, B). In both cell lines, expression of the mutant H3.3 transgene resulted in a significant increase in missegregation events (lagging or bridged chromosomes in the spindle midzone), compared to cells expressing H3.3^WT^-GFP. To test whether these chromosome segregation errors were dependent on levels of phospho-Ser31 rather than mutant transgene expression, we expressed a double mutant containing the K27M driver mutation and a phosphomimetic S31E. Here the inclusion of the phosphomimetic along with the K27M mutation leads to significantly lower frequency of chromosome missegregation in both cell types, suggesting the effect of the K27M on chromosome segregation is dependent on lowering Ser31 phosphorylation levels, which are rescued by the phosphomimetic (Figure 2). We further confirmed these transgene results using the SF-9427 pediatric GBM cell line that contains wild-type histone H3 (Supplemental Figure 2). Expression of either K27M or S31A significantly increased the frequency of chromosome missegregation events in SF-9427 cells, while expression of the K27M/S31E double mutant suppressed chromosome missegregation.

**Figure 2.**
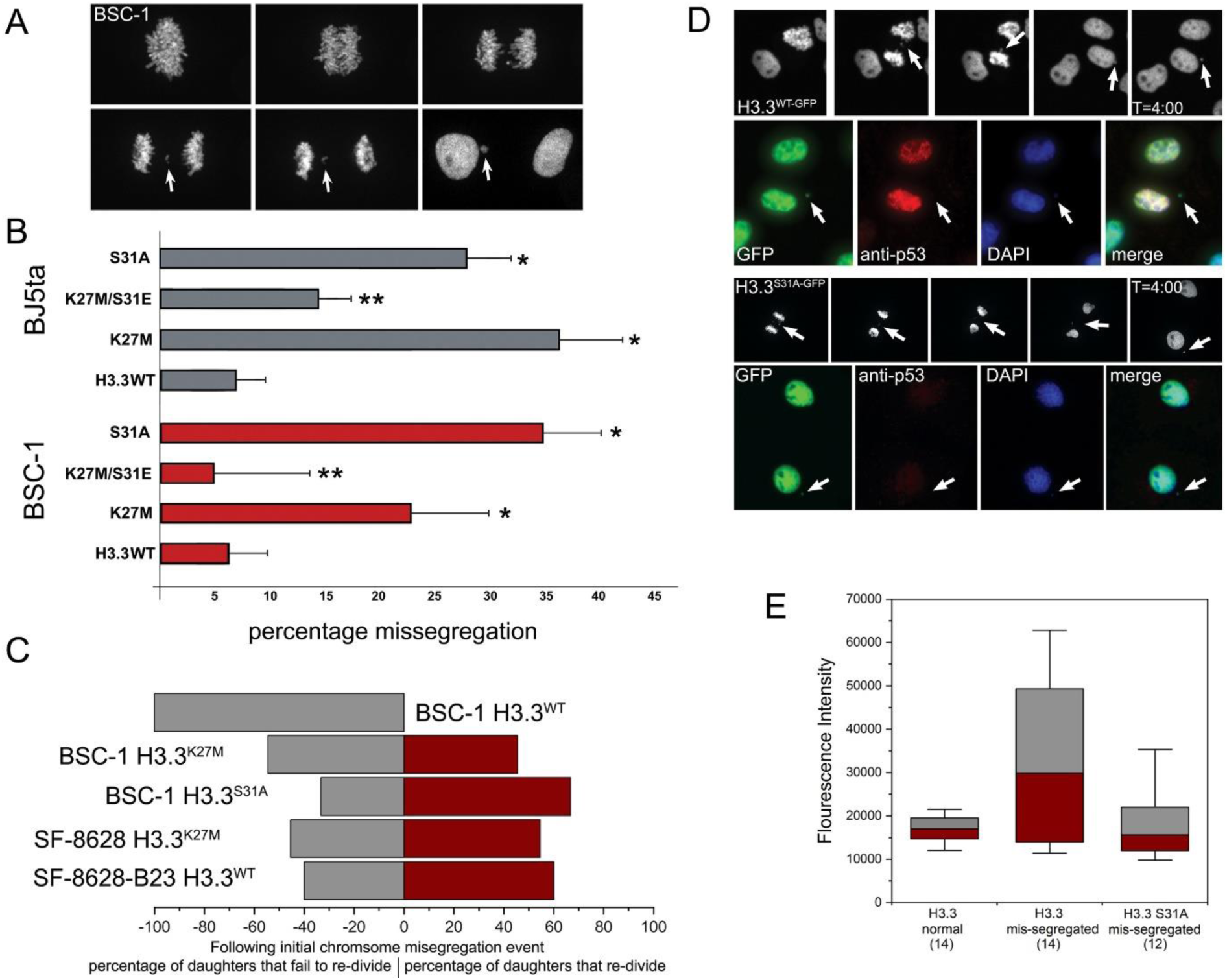
Altering Ser31 phosphorylation in non-transformed cells induces chromosome missegregation and loss of cell cycle checkpoint control. **A**. Live-cell fluorescence of chromosome missegregation in a BSC-1 cell expressing H3.3^K27M^-GFP. **B**. Percentage of anaphase cells with lagging or bridged chromosomes in response to expression of H3.3-GFP constructs in BJ5ta and BSC-1 cells. *n* = 50 cells for each construct. Graph shows mean from five separate experiments ± SD. \**P* < 0.05, \*\**P* ≥ 0.05, two-tailed *t*-tests compared to H3.3WT expressing cells. **C**. Bar graph showing cell cycle fate of daughter cells derived from an initial mitotic division where one or more chromosomes missegregated. Compares percentage of daughter cells that undergo a second mitotic division to the percentage of those cells that fail to divide a second time. Because BSC-1H3^WT^ cells do not spontaneously missegregate chromosomes, all cells were subjected to chilling/re-warming, followed by live-cell imaging. SF-8628 and SF-8628-B23 cells expressed H2B-GFP as a chromosome live-cell marker, BSC-1 cells expressed requisite H3.3-GFP construct that act as both mutant and chromosome marker. *n* = 10 cells for BSC-1^H3.3WT-GFP^, 6 cells for BSC-1^H3K27M-GFP^, 6 cells for BSC-1^H3S31A-GFP^, 11 cells for SF-9628^H3K27M^, and 10 cells for SF-9628^H3WT^. Daughter cell number is 2x original cell number. One of the BSC-1^H3K27M^ daughter cells was lost from the experiment, giving an odd number of daughters analyzed. **D**. Live-cell/fixed cell analysis of post-mitotic p53 expression following chromosome missegregation for H3.3^WT-GFP^ vs H3.3^S31A-GFP^ expressing cells. Following chromosome missegregation (arrows), cells were imaged for four hours, position of the cell marked with a diamond scribe in the microscope nosepiece, and the coverslip fixed and immuno-labelled with anti-p53. **E**. Box and Whisker plots (median value, 50^th^ and 75^th^ percentile, Max and Min) of p53 levels, as measured from post-mitotic cells in **D**. The levels of nuclear p53 after four hours in cells with a missegregated chromosome are compared to nuclear p53 levels after four hours in BSC1 cells that divide normally. *n* = 14 daughter cells for BSC-1^H3.3WT-GFP^ that divide normally, 14 daughter cells for BSC-1^H3.3WT-GFP^ that missegregate a chromosome, and 12 daughter cells for BSC-1^H3.3S31A-GFP^ that missegregate a chromosome. Each is an independent experiment.

Many human tumors, including DIPGs, are aneuploid, and established tumors often settle on characteristic karyotypes that balances numerical chromosomal abnormalities with compensating mutations in tumor suppressors and other signaling genes. However, it should be noted that a direct experimental link between chromosome instability and tumorigenesis remains elusive and can be cell and/or tissue type specific^28-31^. In many cases – but not all – chromosome segregation defects can induce the stabilization and accumulation of nuclear p53, resulting in cell cycle arrest, which can be either dependent or independent on the DNA damage response^10, 27^. In other cases, loss of cell proliferative capacity is dependent on the cGAS-STING pathway that monitors dsDNA in the cytoplasm; the missegregated chromosome forms a micronucleus in G_1_ that is often leaky, and resulting in dsDNA interacting with the cytoplasm, which triggers STING activation and senescence ^30, 31^. However, tumor cells often bypass both the cell cycle arrest and the cellular senescence mechanisms in order to proliferate as chromosomally unstable clones^31^.

To examine the role of H3.3 driver mutations in cell cycle arrest in non-transformed cells we analyzed cell cycle progression following chromosome missegregation in BSC1 cells transiently expressing H3.3^WT^, H3.3^S31A^, or H3.3^K27M^. We previously demonstrated that BSC1 cells arrest in G_1_ in response to chromosome missegregation^10^. Importantly, the expression of either the S31A or K27M mutation induces chromosome missegregation on their own, while the H3.3^WT^ cells do not. Thus, missegregation must be experimentally induced by chilling-rewarming, which in itself does not alter cell viability of cell cycle progression, as previously shown^10, 32^.

At three-days post-transfection, cells on coverslips were assembled into imaging chambers, individual cells identified in metaphase using spinning disk confocal microscope, and imaged as they progressed into anaphase. If one or more chromosomes missegregated, the daughter cells were followed by time-lapse microscopy for 70 hrs. Consistent with our previous results we did not observe any H3.3^WT^-expressing cells that missegregate a chromosome subsequently undergoing a second round of division – both daughters remained arrested for the duration of our observations (Figure 2C and Supplemental Figure 3). However, for both the K27M and S31A-expressing cells, approximately 50% of the daughters resulting from an anaphase with a chromosome missegregation underwent a second division, suggesting these nascent aneuploid cells have a compromised cell cycle checkpoint function – the definition of chromosome instability (Figure 2C).

**Figure 3.**
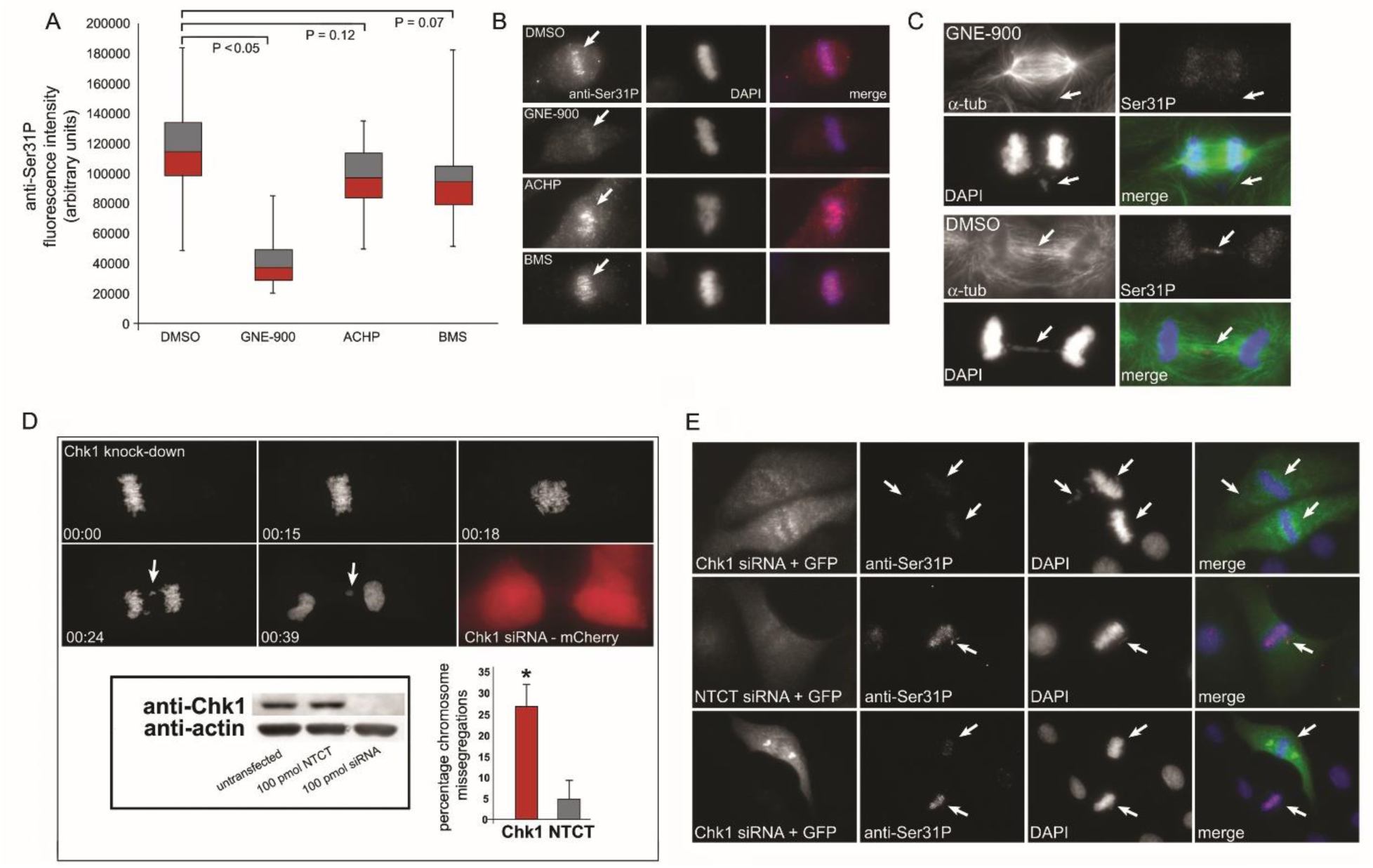
Chk1 kinase phosphorylates H3.3 Ser31 during mitosis. **A**. Box and Whisker plots (median value, 50^th^ and 75^th^ percentile, Max and Min) of pericentromeric phospho-Ser31 levels of mitotic BJ5ta cells treated for 1 hour with either a Chk1 inhibitor (GNE-900; 10µM), or one of two IKKα inhibitors (BMS; 10 µM) or (ACHP; 10 µM). *p* values are from two-tailed *t*-tests. **B**. Metaphase cells from **A**, showing levels of pericentromeric phospho-Ser31 (arrows). **C**. Phospho-Ser31 on missegregated chromosomes (arrows) in post-anaphase cells treated with GNE-900 or DMSO. Phospho-Ser31 is lost from lagging chromosomes in the GNE-900 treated cells (arrow). **D**. Frames from a live-cell imaging sequence of an H2B-GFP expressing BSC-1 cell nucleofected with Chk1 siRNA + mCherry fluorescent protein. The cell missegregates a chromosome (arrow). Anti-Chk1 immunoblot demonstrating siRNA knock-down compared to non-targeted control siRNA. Bar graph of percentage of chromosome missegregations in Chk1 siRNA vs NTCT siRNA ± SD. *n* = 30 cells per condition from three separate experiments. \**P* < 0.05, two-tailed *t*-test compared to NTCT siRNA transfected cells. **E**. Comparison of phospho-Ser31 labelling in mitotic BSC-1 cells nucleofected with Chk1 siRNA + EGFP fluorescent protein vs NTCT siRNA. In the bottom panels note the difference in levels of phospho-Ser31 (arrows) in two adjacent mitotic cells, one expressing Chk1 siRNA (green) and the other un-transfected.

We also tested whether K27M-mutant DIPG cells (SF-8628) can undergo repeated rounds of division following chromosome missegregation and found that ∼50% of K27M mutant daughter cells underwent one or more subsequent cell divisions (Figure 2C). However, when we analyzed SF-8628-B23 cells using the same assay, we found that reversion of the H3.3 K27M to H3.3 wildtype was not sufficient to restore G_1_ arrest. This is not overly surprising, given that tumor cells obtained from DIPG patients accumulate extensive secondary mutations in addition to the H3.3 driver mutations. From these data we conclude that expression of H3.3 mutants K27M or the non-phosphorylatable version S31A interferes with the ability of cells to respond to checkpoint signaling following chromosome missegregation. Further defining the checkpoint mechanisms that couple chromosome missegregation to the cell cycle machinery is necessary to determine the role played by Ser31 phosphorylation.

We previously used single cell analyses to demonstrate that in BSC1, cells accumulate p53 following chromosome missegregation and this is dependent on H3.3 Ser31 phosphorylation. Here we expressed H3.3 S31A-GFP in BSC1 cells, identified an individual cell undergoing chromosome missegregation by live-cell imaging, and followed that cell for 4 hours post anaphase, a time when p53 is known to accumulate to maximal levels in this cell line. The cell was then fixed and labeled with anti-p53 (Figure 2D). We compared post-anaphase p53 levels in these S31A-expressing cells to that of BSC1 cells expressing H3.3 WT-GFP. H3.3^WT^-expressing cells that divide all chromosomes normally have basal levels of nuclear p53, while those WT-expressing cells that missegregate a single chromosome accumulate significant levels of p53 in their post-anaphase nuclei (Figure 2E). In contrast, H3.3^S31A^-expressing cells that missegregate a single chromosome do not accumulate nuclear p53 (Figure 2E), suggesting that decreased phospho-Ser31 inhibits the induction of the p53-dependant aneuploidy checkpoint/fail-safe response to chromosome in G_1_.

Histone H3 residues Ser10, Ser28, and Ser31 are each phosphorylated by multiple kinases, and the kinase specificity appears to depend upon cell cycle stage^2, 33^. For example, Aurora B kinase phosphorylates H3 Ser28 as cells enter mitosis, whereas during interphase Ser28 phosphorylation is catalyzed by MSK1/2 kinase in response to MEK/ERK/p38 signaling^2^. In addition to cell cycle regulation, the distribution of phosphorylated histone H3 sub-species is also thought to be region specific^1,2^. In the case of mitotic H3.3, Ser31 phosphorylation is normally restricted to heterochromatin located in the pericentromere (Figure 3A). However, phospho-Ser31 can spread to chromosome arms on chromatids that become mispositioned^10^, or in cells with the Alternative Lengthening of Telomeres (Alt) phenotype^3^. The mechanisms underlying this positional specificity are not known. The identity of the mitotic Ser31 kinase also remains controversial. Previous studies have identified Serine/threonine kinase Chk1 as a potential mitotic Ser31 kinase, but certain Chk1 inhibitors failed to block Ser31 phosphorylation on lagging chromosomes or even at the pericentromere^3, 5, 10^. Adding to this, a recent study demonstrated that during interphase histone H3.3 deposited in regions of active transcription undergo Ser31 phosphorylation by IKKα kinase, and this phosphorylation is required to amplify stimulation-induced gene transcription^6^. In order to further clarify the identity of the mitotic kinase of H3.3 Ser31, we treated BJ5ta cells with an inhibitor to Chk1 (GNE-900) or two different inhibitors to IKKα kinase (ACHP or BMS) and measured Ser31 phosphorylation levels in individual mitotic cells by fluorescence microscopy. GNE-900 treatment resulted in a significant decrease in mitotic Ser31 phosphorylation, while neither of the IKKα kinase inhibitors did (Figure 3A, B). Treatment with GNE-900 also decreased phospho-Ser31 on chromosomes lagging in the spindle midzone (Figure 3C). We found similar results using a different Chk1 inhibitor AZD7762 (Supplemental Figure 5C, D).

To confirm that Chk1 acts as a mitotic Ser31 kinase, we used previously validated siRNAs to knock-down expression of Chk1 (*CHEK1*) and tested whether loss of this kinase reduced Ser31 phosphorylation levels and whether this loss of phospho-Ser31 resulted in an increase in chromosome missegregation^33-34^. Transfection with 100 pmol of Chk1 siRNA resulted in loss of detectable Chk1 protein, and live-cell imaging of Chk1 knock-down cells revealed a significant increase in chromosome missegregation events compared to cells transfected with a non-target control siRNA (Figure 3D). This is consistent with previous work, showing Chk1 plays a key role in maintaining chromosome stability, in addition to its canonical function in DNA damage response^34-36^. Chk1 siRNA also abolished mitotic Ser31P labelling of both pericentromeric regions and labelling of chromosomes that fail to congress at metaphase – specificity for the effects of the Chk1 siRNA is controlled for by the presence of adjacent, untransfected mitotic cells in the same preparation, which retain their mitotic anti-phospho-Ser31 labelling (Figure 3E).

We next tested whether Chk1 can directly phosphorylate H3.3 protein using an in vitro kinase assay, with recombinant H3.3 protein as the substrate and purified, active Chk1 kinase as the enzyme. Chk1 is known to phosphorylate all H3 variants at the highly conserved Thr11 residue^37^ and phospho-Thr11 acted as a positive control for our assay. Chk1 phosphorylated full length H3.3 protein in vitro, and the introduction of the S31A mutation reduced Chk1 phosphorylation but did not abolish it (Supplemental Figure 5A). When we analyzed the T11A/S31A double mutant, we found loss of Thr11 and Ser31 together completely abolished detectable Chk1 phosphorylation of H3.3 in vitro.

We further examined Chk1 phosphorylation in vitro using the H3.3 driver mutant K27M as a substrate. To analyze the kinase reactions of single mutant H3.3 proteins (K27M), we used anti-phospho-Ser31 immunoblots (Supplemental Figure 5B). Compared to WT H3.3, the K27M mutation caused an ∼50% reduction in Ser31 phosphorylation, whereas the S31A mutant completely abolished Ser31 phosphorylation. This suggests that the presence of mutations in the histone proteins themselves alters the ability of the Ser31 residue to be phosphorylated. These data further confirm the idea that loss of phospho-Ser31 results in defects in chromosome segregation, and that the presence of the K27M mutation lowers Ser31 phosphorylation levels.

While the presence of H3.3K27M mutation alone is not sufficient to drive gliomagenesis in mice, whereas the addition of platelet-derived growth factor beta (PDGFβ) along with mutant histone expression has been shown to induce brainstem gliomas in murine models that mimic human diffuse midline gliomas^14, 18, 38^. To investigate whether the presence of the S31A non-phosphorylatable H3.3 can also serve to drive these gliomas, we generated a construct where PDGFβ and H3.3 are linked in the same RCAS viral vector with P2A sequence (Supplemental Figure 6). This ensured that all cells that all infected express both exogenous PDGFβ and H3.3. We compared survival of Nestin-TVA mice injected at birth with either PDGFβ-P2A-H3.3^WT^, PDGFβ-P2A-H3.3^K27M^, or PDGFβ-P2A-H3.3^S31A^ (Figure 4A). 46% of the H3.3^WT^ mice generated diffuse low-grade HA-positive tumors, while the remainder were tumor-free, and none of this cohort died over the 100-day experiment. Both the H3.3^K27M^ and H3.3^S31A^ resulted in significant generation of tumors that histologically resemble high-grade gliomas (Figure 4B). Both are positive for Nestin and Ki-67, suggesting they remain in a pluripotent state, and that they are proliferative (Figure 4B). Expression of these H3.3 mutants resulted in a significant decrease in overall survival, compared to the H3.3^WT^ (*p* < 0.05). No significant difference was observed between the survival of mice injected with H3.3^K27M^ and those injected with H3.3^S31A^ (*p* > 0.05).

**Figure 4.**
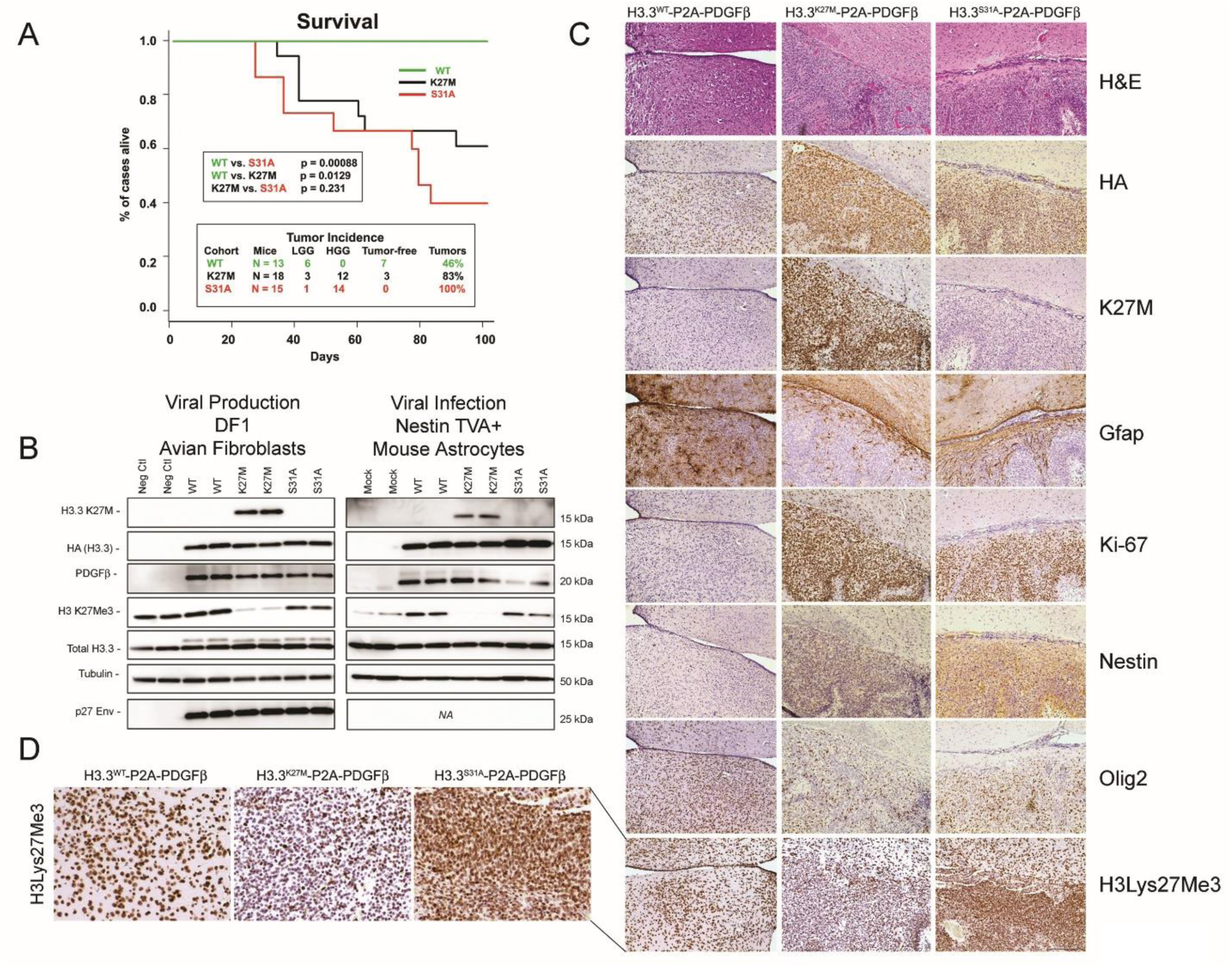
H3.3^S31A^ reduces survival and promotes the development of high-grade gliomas in a genetic mouse model. **A**. Kaplan-Meier percent survival curve of Nestin-TVA mice injected at birth with either PDGBβ-P2A-H3.3^WT^ (green line *n* = 13), PDGBβ-P2A-H3.3^K27M^ (black line *n* = 18) or PDGBβ-P2A-H3.3^S31A^ (red line *n* = 15). A significant difference was observed between the survival of mice injected with H3.3^wt^ containing viruses and H3.3^K27M^ (*p* = 0.0129) and H3.3^S31A^ containing viruses (*p* = 8.8 ×10^−4^). No significant difference was also observed between the survival of mice injected with H3.3^K27M^ and H3.3^S31A^ (*p* = 0.231). **B**. Western blot analysis of DF1 viral producing cells infected with RCASBP(A) PDGBβ-P2A H3.3^WT^, PDGBβ- P2A H3.3^K27M^, and PDGBβ-P2A H3.3^S31A^. The blots were probed with either anti-HA antibody to detect H3.3(-HA), PDGBβ antibody to detect the expression of PDGBβ, H3.3^K27M^ antibody to detect the presence of the H3.3 lysine 27 to methionine mutation and antibodies specific for H3 lysine 27 tri-methylation. Blots were probed with P27 antibody to detect the presence of the RCAS(A) viral envelope and H3.3 and β-tubulin antibodies to confirm equal loading. Western blot analysis of Nestin-TVA+ primary astrocytes infected with viruses containing RCASBP (A) PDGBβ-P2A H3.3^WT^, PDGBβ-P2A H3.3^K27M^, and PDGBβ-P2A H3.3^S31A^. The blots were probed with either HA antibody to detect virally delivered H3.3(-HA), PDGBβ antibody to detect the expression of PDGBβ, H3.3^K27M^ antibody to detect the presence of the H3.3 lysine 27 to methionine mutation, and antibodies specific for H3 lysine 27 tri-methylation, which confirmed a reduction in global in the H3.3 lysine 27 to methionine mutant cells. Blots were probed with β-tubulin and anti-H3.3 antibody to confirm equal loading. **C**. Representative hematoxylin and eosin (H&E) staining and immunohistochemistry (IHC) on serial sections of mouse brains for a HA epitope tag on virally delivered H3.3 confirmed H3.3^WT^, H3.3^K27M^ and H3.3^S31A^ expression and allowed tumor grade and the percentage of mice with tumors to be estimated. While none of the Nestin-TVA mice injected with PDGBβ-P2A H3.3^WT^ died within 100 days 6/13 (46%) were found to have areas of diffuse HA nuclear positive tumor cells, 15/18 (83%) of PDGBβ-P2A H3.3^K27M^ injected mice and 15/15 of 100% of PDGBβ- P2A H3.3^S31A^ injected mice were found to have brain tumors. The majority 14/15 (93%) of the H3.3^S31A^ and 12/18 (67%) of the H3.3^K27M^ tumors histologically resembled high-grade gliomas; they possessed well-demarcated necrosis, pronounced aberrant angiogenesis, pleomorphic nuclei, and were diffusely infiltrative into the brain parenchyma. IHC demonstrated that all tumor cells strongly express oligodendrocyte differentiation marker Olig2. Expression of astrocyte differentiation marker Gfap was present in reactive astrocytes within the tumor but not a clear feature of the tumor’s cells. None of the H3.3^WT^ tumor cells expressed the stem cell marker Nestin; however, both the high-grade H3.3^S31A^ and H3.3^K27M^ tumors expressed Nestin, suggesting that they remain in a pluripotent state. Assessment of cellular proliferation using IHC for the proliferation marker Ki67 demonstrated that the high-grade H3.3^S31A^ and H3.3^K27M^ tumors were highly proliferative, whereas mitotic figures were extremely rare in low-grade H3.3^WT^ tumors. IHC with antibodies specific for the H3.3 lysine 27 to methionine mutations confirmed H3.3^K27M^ expression in the tumor cells. IHC and antibodies specific for H3 lysine 27 tri-methylation confirmed a reduction in global H3 methylation in H3.3^K27M^ tumors compared to H3.3^WT^ tumors and normal brain cells. H3 lysine 27 tri-methylation was not reduced in H3.3^S31A^ tumors compared to H3.3^WT^ tumors and normal brain cells. **D**. A magnified view of IHC on cells in the central tumor region stained for H3 lysine 27 tri-methylation shows reduction in triple methylation levels in the K27M tumors compared to WT or S31A. Scale bars represent 200 µm.

We examined tumor tissue slices from mice using the anti-K27M antisera: cells from the H3.3K27M mice were positive and those from the H3.3WT and H3.3S31A mice were negative for K27M. Also, we found robust labelling with anti-H3Lys27Me3 antisera in tissue slices from the H3.3WT and H3.3S31A mice, as well as avian fibroblasts and TVA+ mouse astrocytes infected with viruses expressing either of these constructs (Figure 4 B). Not surprisingly, H3Lys27Me3 labelling was significantly decreased in both cells and tissue from the H3/3K27M mice (Figure 4B, D) consistent from previous reports that the presence of H3.3 K27M induces a decrease in Lys37 triple methylation^13, 14, 18^. We found similar results analyzing either normal cells expressing exogenous H3.3Ser31A-GFP or our DIPG cells CRISPR-edited to express H3.3 S31A (Supplemental Figure 7). This observation is extremely interesting, because the mechanism thought to underly the action of oncogenic histone mutations in the induction of diffuse midline gliomas is solely the loss of H3Lys27 triple methylation leading directly to epigenetic dysregulation. Here our results suggest that the formation of diffuse glial tumors in mice that closely mimic K27M gliomas does not require the loss of H3Lys27 methylation.

We also examined the levels of H3.3 phospho-Ser31 in mitotic mouse TVA+ astrocytes infected with RCAS virus expressing H3.3^WT^-GFP, H3.3^K27M^-GFP, or H3.3^S31A^-GFP by immunoblot (Supplemental Figure 8). Mitotic cells were enriched by nocodazole treatment and mechanical shake-off. Not surprisingly, Ser31 phosphorylation is lost from the H3.3S31A non-phosphorylatable mutants but the K27M mutant H3.3 protein has decreased capacity for mitotic Ser31 phosphorylation, consistent with our findings from cultured cells and in vitro kinase assays.

Together our results reveal that the ability of histone H3.3 protein to become phosphorylated at position Ser31 during mitosis is altered by the presence of the K27M mutation. The decreased phosphorylation results in an increase in missegregated chromosomes during anaphase that lack sufficient phospho-Ser31 on their arms to activate the p53 response to aneuploidy. By both generating missegregated chromosomes and short-circuiting the cell cycle response to the aneuploidy, the H3.3 K27M mutation appears to drive chromosome instability, in addition to its dominant-negative effect on epigenetic dysregulation^13, 14, 18, 19^. This suggests that the downregulation of Ser31 phosphorylation by H3.3 mutations in early gliomagenesis results in a population of cycling cells with aneuploid karyotypes. The early onset of aneuploidy combines with the extensive epigenetic re-programming driven by the K27M effect of Lys27 triple methylation. The net result is gliomagenesis That nearly all gliomas are aneuploid highlights this convergent tumor evolution. Our work suggests that these signature histone H3.3 mutations may be particularity oncogenic because they provide a double hit the induces both epigenetic dysregulation and chromosome instability.

## Acknowledgments

This work supported by grants from the Department of Defense (CDMRP CA171071 and CA200747 to EHH), the Minnesota Partnership for Biotechnology and Medical Genomics (to DJD, JPR, and EHH), the National Institutes of Health (R01NS117432 to DJD), the American Cancer Society (Research Scholar Award to JPR), and by the Hormel Foundation. CAD was supported by the National Cancer Institute T32 CA21783602 Neuro-Oncology Training Grant to the Mayo Clinic. We thank the Marit Mary Swenson Charitable Trust for their support and dedicate this work in her memory.

