## Supplemental Data for "The histone H3.3 K27M mutation found in diffuse midline gliomas coordinately disrupts adjacent H3.3 Ser31 phosphorylation and the fidelity of chromosome segregation"

**Supplemental Material (Day et al., 2022)**

**Supplemental Figures 1-8**

**Materials and Methods**

**Additional references**

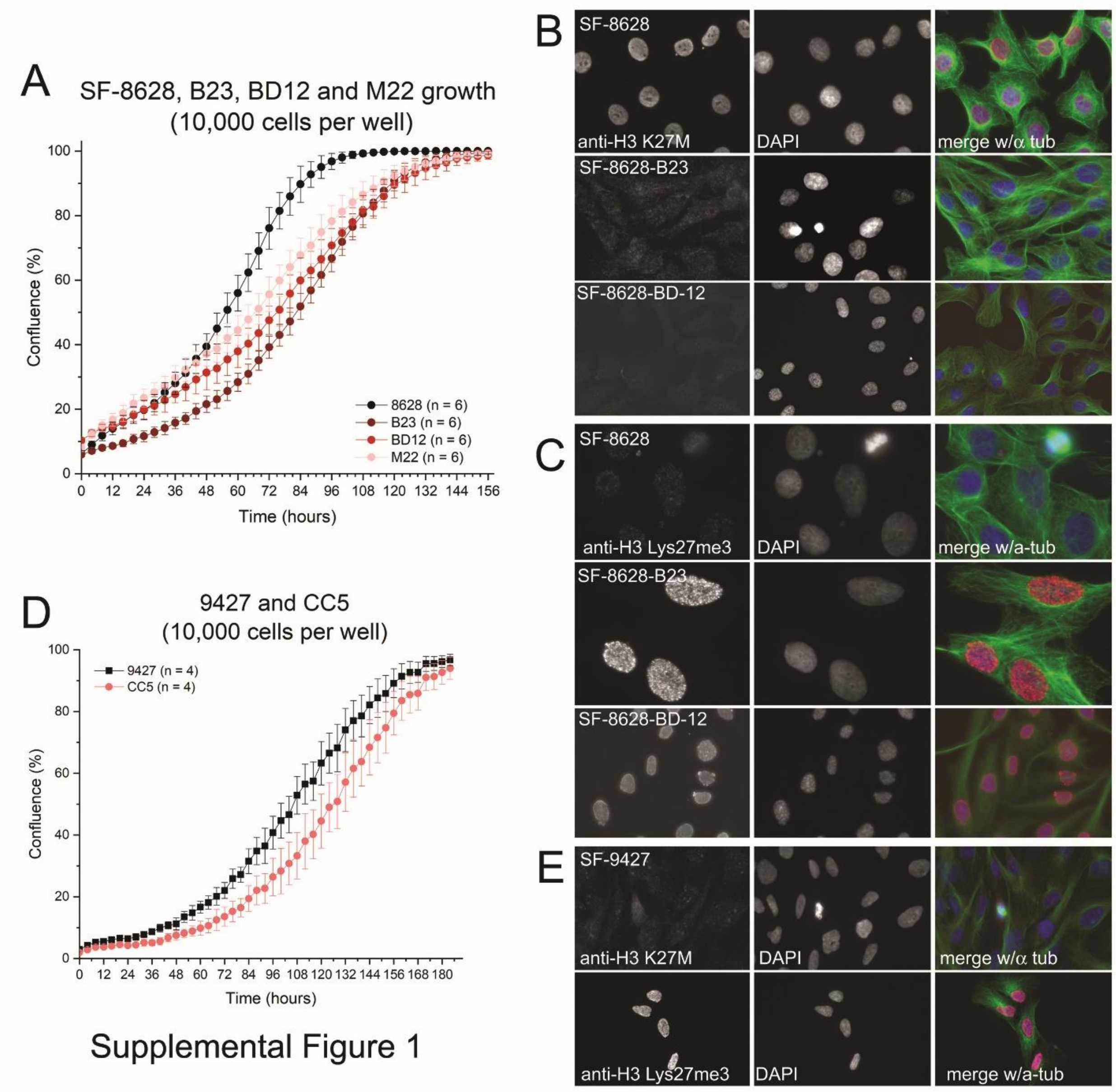


**Supplementary Figure 1. CRISPR gene editing. A.** Live-cell proliferation curves generated with an Incucyte, showing growth of SF-8628$^{H3.3K27M}$ cells and CRISPR-edited SF-8628 cells with either H3.3$^{M27K}$ reversion (B23 and BD12) or H3.3$^{M27K}$ reversion + S31A non-phosphorylatable (M22). The time to reach 100% confluence in the B23, BD12 and M22 cells (t = 168 hrs) lags slightly behind the parental SF-8628 cells (t = 126 hrs). **B.** Interphase cells labelled with anti-K27M detects the nuclear K27M mutant H3 protein in SF-8628 cells; this is lost from nuclei in the revertant B23 and BD12 cell lines. The SF-8628 cells have significantly decreased H3Lys27Me3 in their nuclei, compared to the B23/BD12 revertant cells. SF-9427, which have $H3^{WT}$ have high levels of H3Lys27Me3 in their nuclei and lack nuclear anti-K27M labelling. **C.** Live-cell proliferation curves generated with an Incucyte, showing growth of SF-$9427^{H3WT}$cells and CRISPR-edited SF-9427 cells with H3.3 S31A non-phosphorylatable (CC5). The time to reach 100% confluence in the CC5 and parental SF-9427 cells is similar.

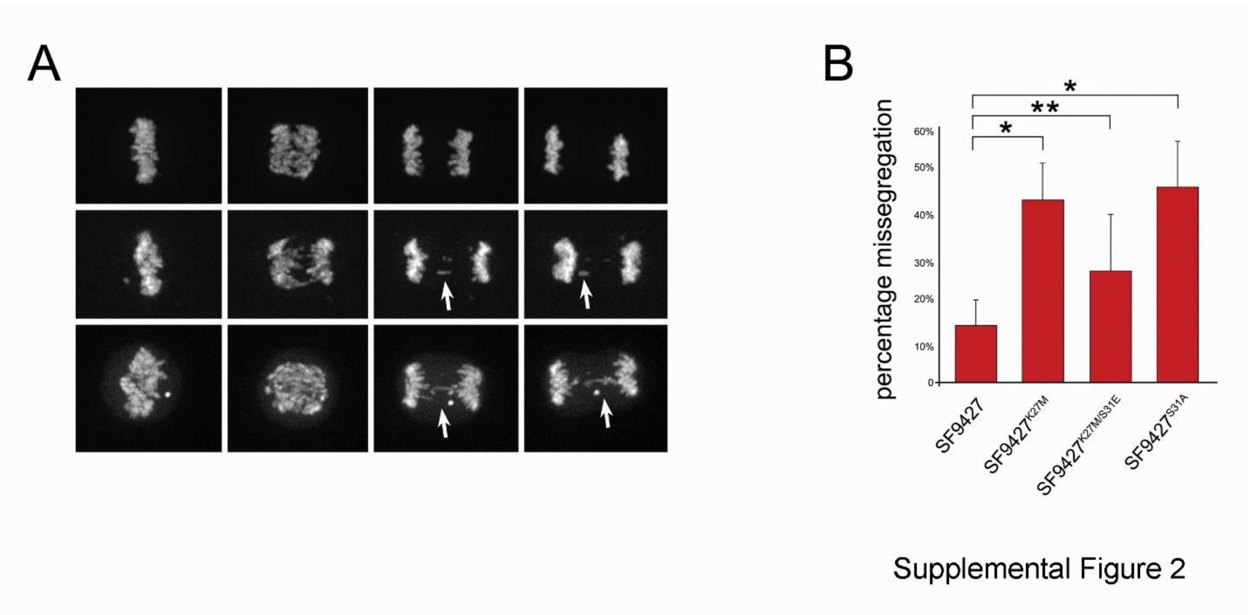


**Supplementary Figure 2. Expression of mutant H3.3 constructs in SF-9427 cells induces chromosome missegregation. A.** Frames from live-cell imaging sequences of SF-9427-H3.3K27M-GFP expressing cells. Chromosome missegregation (arrows) resulting in lagging chromosomes or bridged chromosomes is clearly visible. **B.** Percentage of anaphase cells imaged by live-cell spinning disk confocal microscopy with one or more lagging or bridged chromosome. *n* = 50 cells. Graph shows mean from five separate experiments ± SD. $^{*}P < 0.05$, $^{**}P \geq 0.05$ two-tailed *t*-tests.

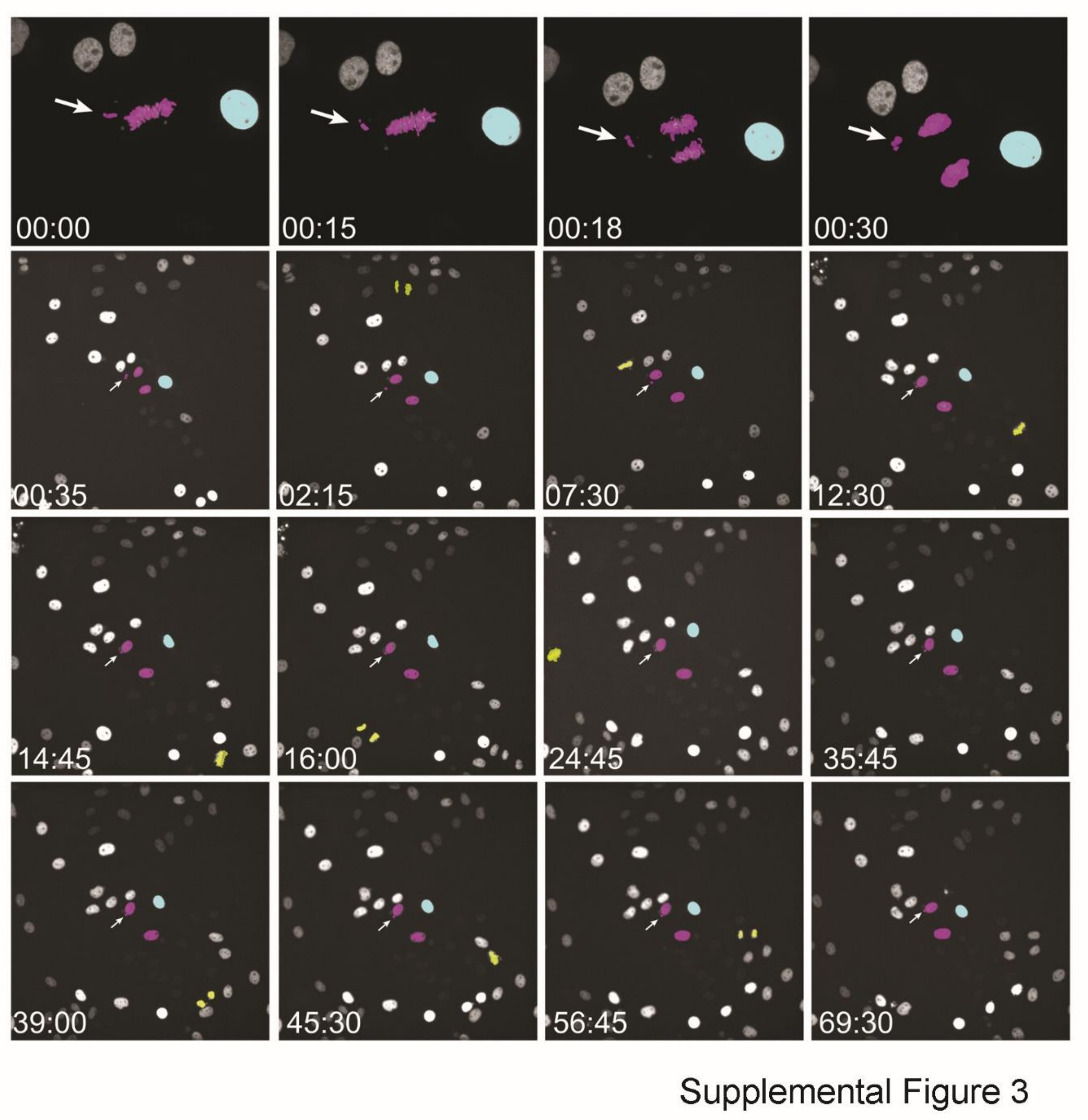


**Supplementary Figure 3. Cell cycle arrest following chromosome missegregation in a normal cell.** Frames from a time-lapse video of a BSC-1 cell expressing H3.3WT-GFP (pseudo-colored magenta) that is induced to missegregate a single chromosome (arrow) in anaphase by chilling re-warming. The daughter cells (one containing the missegregated chromosome as a micronucleus – arrow) do not undergo a second round of cell division for the next ~70 hrs. Other cells in the field of view continue to divide (pseudo-colored yellow). T = Hr:min. Spinning disk confocal imaging.

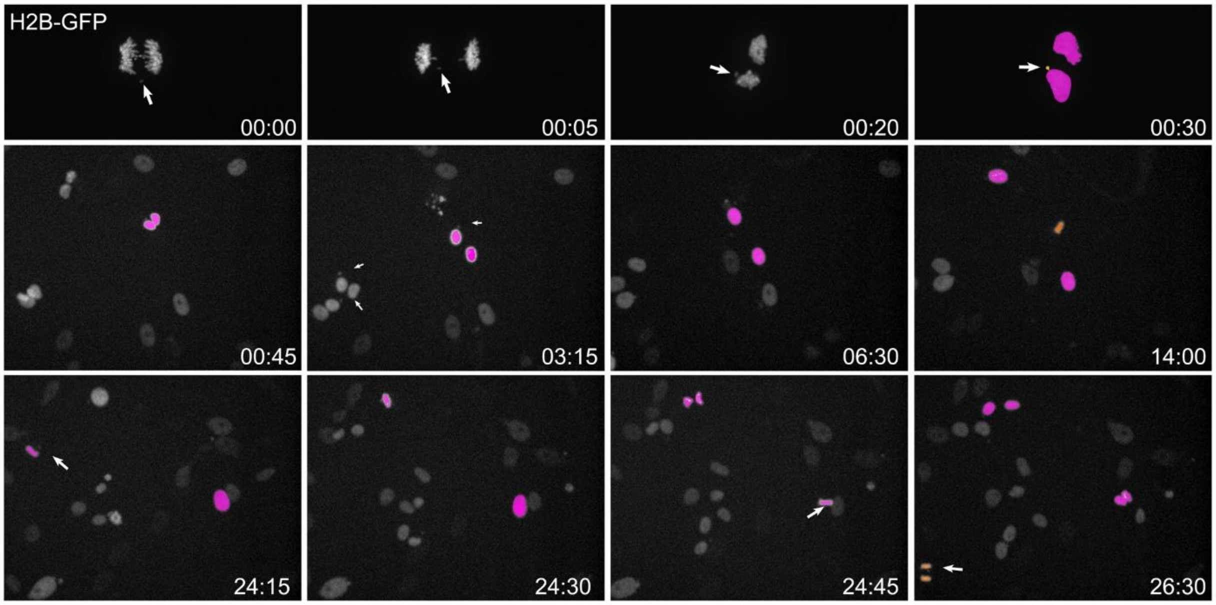


Supplemental Figure 4

**Supplementary Figure 4. Repeated mitotic divisions following chromosome missegregation in K27M-expressing cells.** Frames from a time-lapse video of a SF-8628 cell expressing H2B-GFP (pseudo-colored magenta) that missegregates a chromosome (arrow). The daughter cells (one containing the missegregated chromosome as a micronucleus – arrow) undergo a second round of cell division for the next ~24 hrs. Other cells in the field of view continue to divide as well (pseudo-colored orange), including cells that continue to missegregate chromosomes (arrow in cell at 26:30). T = Hr:min. Spinning disk confocal imaging.

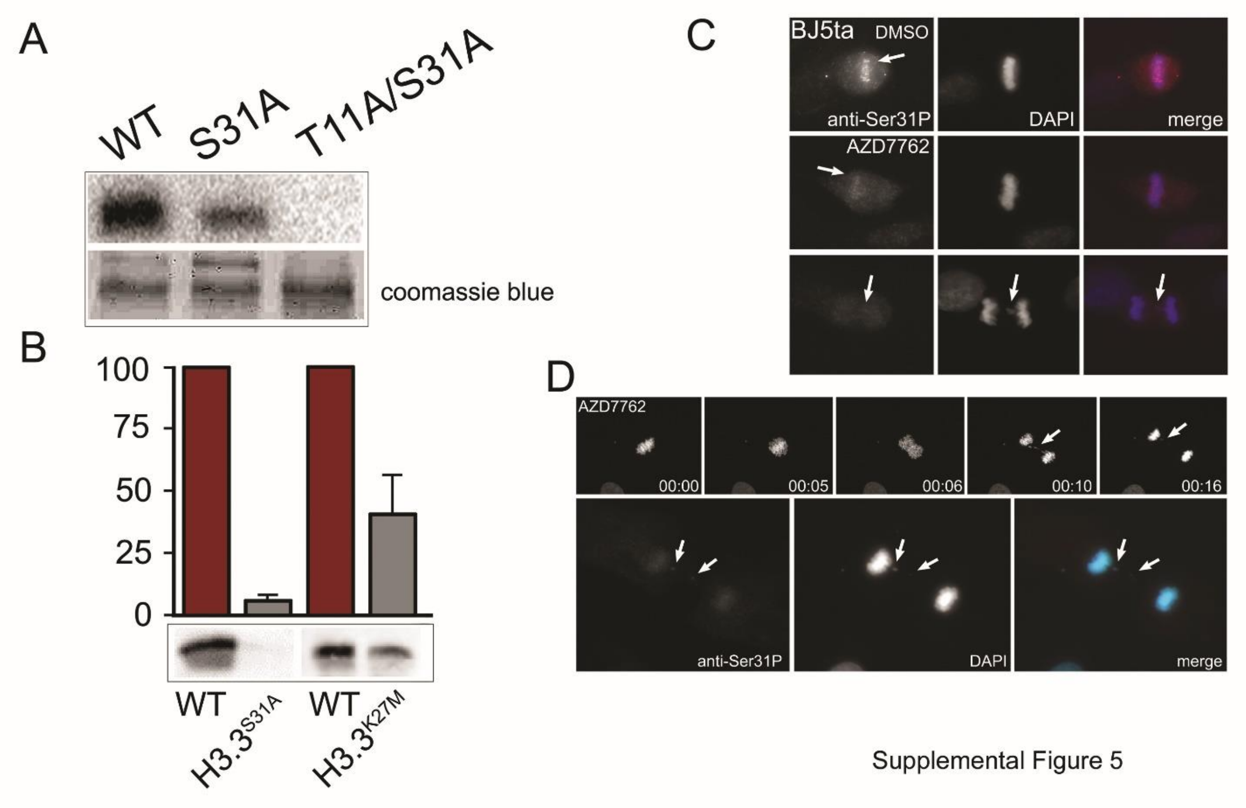


**Supplementary Figure 5. Chk1 kinase phosphorylates H3.3 Ser31 in vitro. A.** In vitro kinase assay using purified Chk1 kinase as the enzyme and purified H3.3 proteins as the substrate. Upper panel: radiograph of $\gamma$-p32-labelled protein, lower panel Coomassie blue labelling of total protein. **B.** Levels of protein phosphorylation of H3.3S31A or H3.3K27M from in vitro kinase assays normalized to phosphorylation of H3.3 WT. Phospho-protein is detected with anti-H3.3Ser31P antibody. Graph shows mean from three separate experiments ± SD. **C.** Mitotic BJ5ta cells treated with DMSO (upper panels) or the Chk1 inhibitor AZD7762 (lower two sets of panels). Inhibition of Chk1 kinase activity results in deceased mitotic H3.3 phospho-Ser31 in metaphase cells and on lagging chromosomes in anaphase cells. **D.** Live-cell/fixed cell analysis of a BJ5ta cell treated with AZD7762. As the cell enters anaphase several chromosomes missegregate (arrow). After fixation and labelling with anti-phospho-Ser31, there are two lagging chromosomes that remain in the central spindle region (arrows). These lack anti-phospho-Ser31 labelling.

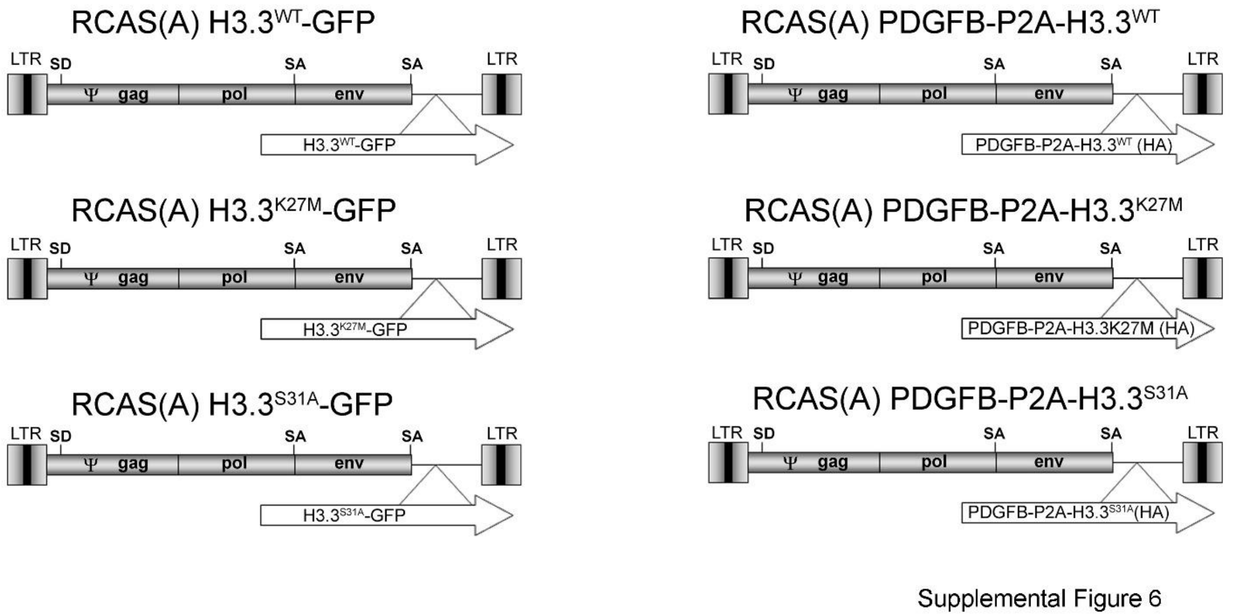


**Supplementary Figure 6. Schematics of RCAS viral vectors used in these studies.**

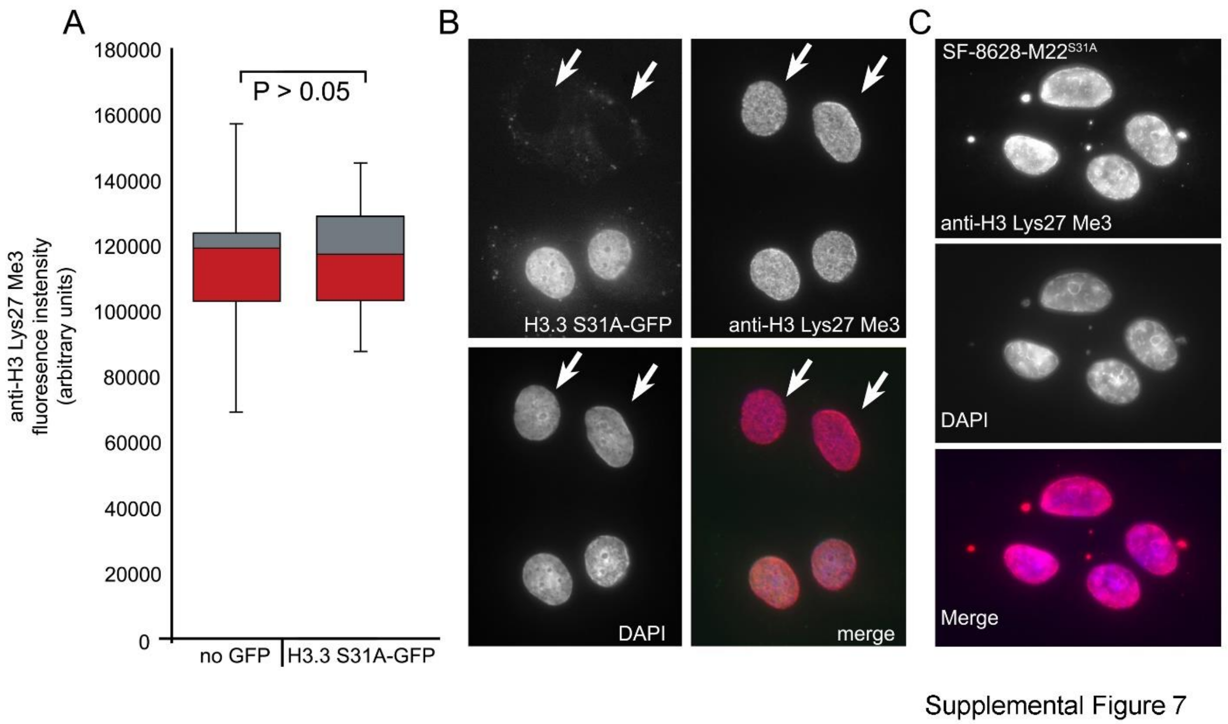


**Supplementary Figure 7. Expression of H3.3S31A does not down-regulate H3Lys27 triple methylation. A.** Box and Whisker plots (median value, 50$^{th}$ and 75$^{th}$ percentile, Max and Min) of nuclear H3Lys27Me3 levels of interphase BJ5ta cells transiently expressing H3.3S31A-GFP, compared to non-expressing cells in the same field of view (examples in **B**). There is no significant difference in the H3Lys27Me3 levels between these two populations of cells. **C.** Interphase SF-8628-M22$^{S31A}$ cells labelled with anti- H3Lys27Me3 antibodies. The nuclei exhibit robust anti- H3Lys27Me3 labelling.

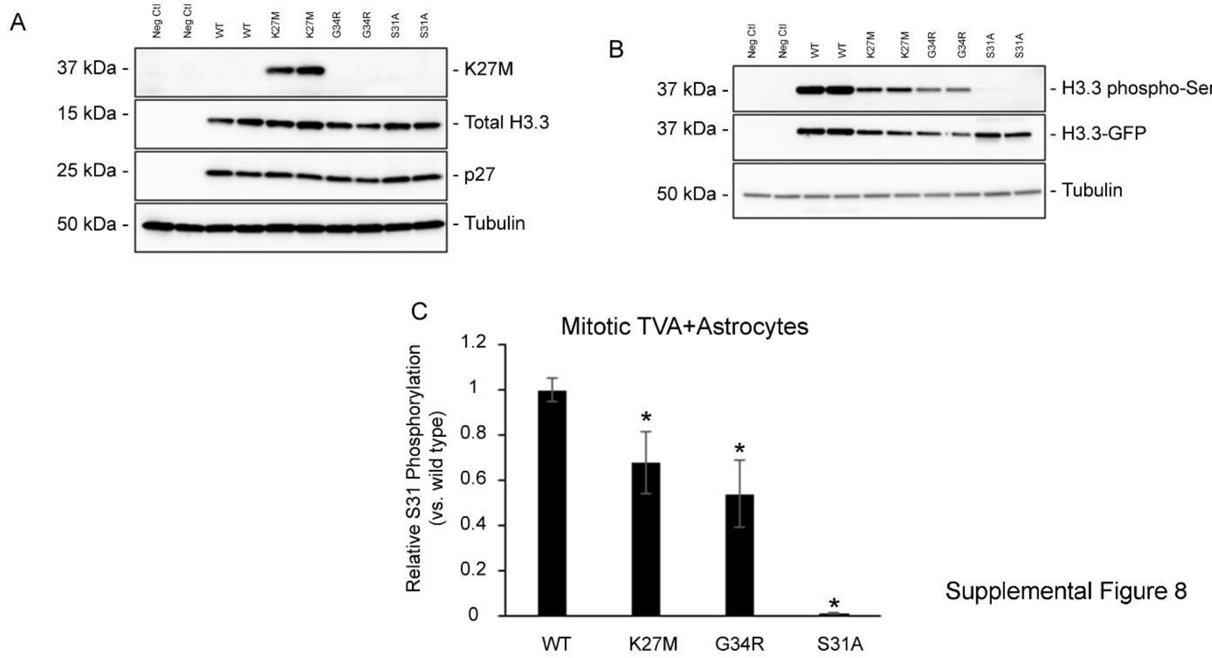


**Supplementary Figure 8. Reduction in H3.3 Serine 31 phosphorylation on H3.3 K27M mutant histones in mouse astrocytes.** In order to determine the level of H3.3 Ser31P on H3.3 K27M mutant histones during mitosis we infected Nestin-TVA+ primary astrocytes with H3.3$^{WT-}$GFP, H3.3$^{K27M}$-GFP, H3.3$^{G34R}$-GFP and H3.3$^{S31A}$ -GFP after flow sorting for GFP expression, confirmation of gene expression and nocodazole synchronization we collected the mitotic faction and performed a western blot analysis of Ser31P on the GFP tagged mutant histones to allow for separation with WT endogenous histone H3.3. **A.** Western blot analysis of DF1 viral producing cells infected with and producing viruses containing RCASBP(A) H3.3$^{WT-}$GFP, H3.3$^{K27M}$-GFP, H3.3$^{G34R}$-GFP and H3.3$^{S31A}$ -GFP (~37kDa). The blots were probed with H3.3 and H3.3$^{K27M}$ antibodies to H3.3 and to detect the presence of the H3.3 lysine 27 to methionine mutation and P27 antibody to detect the presence of the RCASBP(A) viral envelope and β-tubulin to confirm equal loading. **B.** Western blot analysis on the mitotic fraction of Nestin-TVA+ primary astrocytes infected with viruses containing RCASBP (A) H3.3$^{WT-}$GFP, H3.3$^{K27M-}$GFP, H3.3$^{G34R}$-GFP, and H3.3$^{S31A-}$GFP following nocodazole synchronizing. The blots were probed with antibodies for H3.3 and antibodies specific for H3.3 Se31P (ab92628, 1:1000 Abcam). Blots were probed with β-tubulin to confirm equal loading. **C**. Quantification of H3.3 Ser31 phosphorylation expression from Nestin-TVA+ primary astrocytes infected with $H3.3^{WT-}$GFP, $H3.3^{K27M}$-GFP, $H3.3^{G34R}$-GFP and $H3.3^{S31A}$ -GFP. Relative levels of H3.3 GFP Ser31P were calculated by densitometry relative to total H3.3-GFP. Normalization was determined by calculating the densitometry for Ser31P and H3.3 and normalizing to tubulin loading control. Data are presented as mean ± SEM. *$P < 0.05$, two-tailed *t*-test compared to $H3.3^{WT}$ infected cells.

### Materials and methods

**Plasmids.** H2B-GFP plasmid was purchased from Clontech (Mountain View, CA). Emerald-H3.3-N-14 and mCherry2-N1 were a gifts from Michael Davidson (Emerald-H3.3-N-14: Addgene plasmid #54116; http://n2t.net/addgene:54116; RRID: Addgene_54116; mCherry2-N1: Addgene plasmid #54517; http://n2t.net/addgene:54517; RRID: Addgene_54517). As the $1^{st}$ *methionine* on the H3 histone tail is cleaved at a very early posttranslational state, the convention is to name histone methylation and mutations starting from the $2^{nd}$ amino acid position[39]. Following this nomenclature, Emerald-H3.3 K27M, S31A, and K27M/S31E were generated by site-directed mutagenesis using Quikchange II (Agilent; Santa Clara, CA). Pet28a_Human_H3.3 was a gift from Joe Landry (Addgene plasmid # 42632; http://n2t.net/addgene:42632; RRID: Addgene_42632). Plasmids encoding His6x-H3.3 S31A, T11A/S31A, and K27M were generated from the Pet28a_Human_H3.3 plasmid by site-directed mutagenesis using Quikchange II.

The retroviral vectors used in this study were replication-competent avian leukosis virus long terminal repeat, splice acceptor, Bryan polymerase-containing vectors of envelope subgroup A RCASBP(A)[40]. H3.3 is encoded by H3F3A NP_002098.1. The H3.3 CDNA was linked to the mature soluble PDGFβ peptide CDNA[38] with a P2A sequences[41] to ensure linked gene expression (otherwise a histone mutant not required for tumor growth or progression may not be included in a tumor). An HA epitope tag was included on the N-terminal of H3.3 to allow for detection of virally delivered. H3.3. K27M, G34R and S31A mutations were introduced into the H3.3 (HA) CDNA and H3.3-GFP CDNA[26]. PDGFβ-P2A-H3.3$^{WT}$, PDGFβ-P2A-H3.3$^{S31A}$, and PDGFβ-P2A-H3.3$^{K27M}$, H3.3-GFP $^{WT}$ H3.3-GFP$^{K27M}$, and H3.3-GFP$^{S31A}$ were cloned into RCASBP(A) DV destination vector using getaway-compatible sequences and LR Clonase Enzyme II (11791-020 ThermoScientific, Waltham, MA) as described[42] and verified by Sanger sequencing.

**In vitro kinase Assay.** His6x-H3.3 variants encoded on pet28a vectors were grown in Rosetta (DE3)pLysS cells (Novagen, Madison, WI) in 2xTY media at 37°C and 250 rpm. Protein induction was carried out by 5 hours of 300 mM IPTG in 2xTY media at 20°C and 300 rpm. Cells were then lysed and His6x-H3.3 bound to Nickel-chelating resin (Novex/LifeTechnologyCompany, Carlsbad, CA). His6x-H3.3 was eluted from the beads using an Imidazole gradient and dialyzed into 1x kinase buffer (50 mM Tris-HCl (SigmaAldrich), 100 mM EDTA (SigmaAldrich), 150 mM NaCl (ThermoFisher), 100 mM EGTA (SigmaAldrich) at pH 7.5 using 7,000 MWCO SnakeSkin dialysis tubing (ThermoFisher). Purified His6x-H3.3 was mixed with purified recombinant Chk1 (Sigma-Aldrich, St. Louis, MO) and either 10 µmol/L of unlabeled ATP or 10 µCi[γ-$^{32}$P]ATP and incubated for 30 minutes at 30°C in 1x kinase buffer. Samples were boiled and run on SDS-PAGE gels. Radiolabeled phospho-H3.3 was detected by autoradiography and silver staining. Non-radiolabeled phospho-H3.3 was analyzed by Western blotting.

**Mammalian Cell lines.** BSC-1, HCT-116, hTert RPE-1, and hTert BJ5ta were acquired from ATCC (Gaithersburg, Maryland) and grown in Dulbecco's Modified Eagle Medium (D2902, Sigma, St. Louis, MO) with 10% FBS (Cytiva Life Sciences, Marlborough, MA), 1x Amphotericin (Corning, Corning, NY), 1x Penecillin-Streptomycin (Cytiva Life Sciences). The pediatric glioma H3F3A WT cell line SF-9427 was obtained from the UCSF Medical Center. The pediatric glioma H3F3A K27M lines, SF-8628 and SF-7761, were acquired from MilliporeSigma (St. Louis, MO). SF-8628, B23, BD12, M22, SF-9427, and CC5 were maintained in DMEM-high glucose (Sigma Cat. No. D6546), 10% FBS, 2 mM L-Glutamine (EMD Millipore Cat. No. TMS-002-C), 1x Amphotericin, and 1x

Penecillin-Streptomycin. SF-7761 were maintained in ReNcell Neuronal Stem Cell Maintenance Medium (MilliporeSigma) supplemented with 20 ng/ml human recombinant EGF (Fisher Scientific, Waltham, MA) and 20 ng/ml human recombinant FGF-2 (MilliporeSigma). SF-7761 were shifted into Dulbecco's Modified Eagle Medium DMEM-high glucose, 10% FBS, 2 mM L-Glutamine, 20 ng/mL EGF, 20 ng/mL FGF-2, 1x Amphotericin and 1x Penecillin-Streptomycin at least 3 passages before experimentation. SF-8628, B23, BD12, M22, SF-9427, M22, SF-7761, RPE and BJ5ta were grown at 37°C and 5% $CO_2$, while BSC-1 which were grown at 37°C and 10% $CO_2$. Nestin-TVA primary astrocytes were established following dissection of newborn Nestin-TVA mouse brain tissue by physical disruption into single cells using scalpels and 0.25% trypsin (25200-114, Gibco, ThermoScientific, Waltham, MA)[43]. DF-1 cells and N-TVA astrocytes were grown in DMEM-high glucose (11995-065, Gibco) supplemented with 10% FBS (A31606-01, Gibco), 1 × penicillin/streptomycin (Gibco 15140-122). DF-1 cells and N-TVA astrocytes maintained at 39 °C. Viral production was initiated by calcium phosphate transfection of 10ug of proviral retroviral vector DNA into 30% confluent DF-1 cells as previously described[44]. To verify function and infectivity N-TVA cells were seeded in 6-well plates at a density of $5 \times 10^4$ cells/well and were maintained in DMEM with 10% FBS. After the cells attached, the medium was removed and replaced with 1 ml of filtered virus-containing medium in the presence of 8 µg/ml polybrene (Sigma, St. Louis, MO) for 2 hrs at 37°C. The virus was removed and replaced with fresh medium, and the cells were incubated at 37°C[45]. WB was used to confirm gene delivery and expression. For mitotic experiments 40% confluent RCAS-H3.3-GFP infected cells were treated with 2mM thymidine (Sigma Aldrich) for 30 hrs. After thymidine treatment, cells were washed with PBS twice and cells were treated with 1 µM nocodazole (Sigma Aldrich) for 12 hrs. Mitotic cells were collected by shake-off and isolated from the media by centrifuging at 1500 rpm for 15 mins.

**CRISPR/Cas9 editing.** To make SF-8628 B23 and BD12 cells, custom Edit-R crRNA (Dharmacon) targeting the K27M encoding region of human *H3F3A* was complexed to Edit-R tracrRNA (Dharmacon, Cambridge, U.K.) for 2 minutes at 94°C in Duplex Buffer (IDT; Coralville, IA) to generate a cr/tracrRNA. K27M specific cr/tracrRNA, recombinant Streptococcus pyogenes Cas9 (Invitrogen, Waltham, MA) and *H3F3A* WT ssDNA were combined in Basic Glial Cell Nucleofection buffer (Lonza, Basel, Switzerland) and nucleoporated into SF-8628 cells using a Lonza Nucfleofector 2b system. Cells were grown as single cell clones in 96 well plates. Once colonies had formed, cells were screened for the presence of the K27M using immunofluorescence with a rabbit anti-H3 K27M antibody (31-1175-00, 1:10,000, RevMab Biosciences) and Alexa-conjugated secondary antibodies (Molecular Probes, Eugene, OR). Clones showing loss of K27M staining had their *H3F3A* gene PCR amplified with direct PCR using Terra PCR Direct polymerase (Takara, Mountain View, CA), 2x Terra PCR direct buffer with Mg2+ and dNTP (Takara) and the endogenous *H3F3A* primer set. Sanger sequencing was performed with the PCR primers.

To generate SF-8628 M22 and SF-9427 CC5 cells, either *H3F3A* K27M or WT specific Edit-R crRNA was complexed for 2 minutes at 94°C Edit-R tracrRNA in Duplex Buffer to generate K27M or WT specific cr/tracrRNA. Cr/tracrRNA, recombinant Streptococcus pyogenes Cas9 and GFP-H3F3A S31A HDR repair plasmid were mixed in Basic Glial Cell Nucleofection buffer and nucleoporated into cells using a Lonza Nucleofector 2b system. Cells were grown as single cell clones in 96 well plates. Colonies were screened for GFP expression using an IncuCyte S3 system. Both endogenous (i.e. non-GFP, non-CRISPRed) and GFP-H3F3A CRISPRed alleles were PCR amplified using Terra PCR Direct polymerase (Takara, Mountain View, CA), 2x Terra PCR direct buffer with Mg2+ and dNTP (Takara) and the endogenous *H3F3A* primer set or GFP-*H3F3A* primers set, respectively. Sanger sequencing was performed with the endogenous *H3F3A* allele primers. Heterozygosity was confirmed in GFP positive clones by allele specific PCR followed agarose DNA gel.

Guide Sequences:

crRNA: *H3F3A* K27M: 5'- AGAGGGCGCACUCAUGCGAGGUUUUAGAGCUAUGCU GUUUUG-3'

crRNA: *H3F3A* WT: 5'-AGAGGGCGCACUCUUGCGAGGUUUUAGAGCUAUGCUGU UUUG-3'

Homology Directed Repair templates:

ssDNA - *H3F3A* WT: 5'-AGCACCCAGGAAGCAGCTGGCTACAAAAGCAGCTCGCA AGAGTGCGCCCTCTACTGGAGGGGTGAAGAA-3'

H3.3 S31A-GFP HDR plasmid: A custom HDR plasmid encoding the 1210 bp of intron 1 at the 5' end followed by the H3F3A cDNA, a 15 amino acid linked, eGFP cDNA, a LoxP cassette containing IRES and Puromycin resistance, followed by SV40 and then 1195 bp of intron 2 at the 3' end. Control CRISPR were done (data not shown) in which CRISPR was performed with and without Cas9 protein. In the absence of Cas9 no GFP expression was achieved in mammalian cells as the HDR plasmid lack an endogenous promoter sequence.

Primer Sequences:

Endogenous human *H3F3A* allele forward: 5'-GTTTGGTAGTTGCATATGGTGATT-3'

Endogenous human *H3F3A* allele reverse: 5'-ACAAGAGAGACTTTGTCCCATT-3'

GFP-tagged human *H3F3A* allele forward: 5'-TTTTCGAGCGGGAAAGGGGT-3'

GFP-tagged human *H3F3A* allele reverse: 5'-CGCTGAACTTGTGGCCGTTT-3'

**Chk1 Inhibitors.** Cells were treated with GNE-900 (150nM; Selleckchem, Houston, TX), ACHP (20 μM; Tocris, Bristol, UK), BMS 345541 (25 μM; Tocris), or AZD7762 (100 nM; Tocris) for four hours then fixed and immune-labelled.

**Chk1 Knock-down.** Pooled NTCT siRNA (siGENOME Non-targeting siRNA #D-001206-14) and Chk1 siRNA were purchase from Dharmacon. Cells were nucleofected with 100 nmol siRNA in cell lines specific buffers using a Lonza Nucleofector 2b. Cells were then grown for 2 days in

DMEM with 10%FBS at 37°C and 10% $CO_2$ before being fixed for immunofluorescence or lysed for Western blotting.

Chk1 siRNA sequence: 5′-CCACAUGUCCUGAUCAUAU-3'

**Western Blotting.** Cell pellets were resuspended in lysis buffer (5 mM PIPES (pH = 7.2), 5 mM NaCl, 5 mM MgCl2, 1 mM EGTA, 1% NP, 10 mg/ml leupeptin, 10 mg/ml pepstatin, 10 mg/ml chymostatin and 10 mg/ml trypsin inhibitor), mechanically lysed manually with a dounce homogenizer, diluted in 2x Laemmli buffer + DTT and boiled. Concentrations of trichloroacetic acid precipitated protein were determined using the Bio-Rad DC Protein Assay (Bio-Rad, Hercules, California). The proteins were separated on a 10%, 4-20% or 10-20% Tris-glycine gradient polyacrylamide gel, transferred to nitrocellulose or PVDF, and incubated for 1 hr at room temperature in blocking solution (0.05% Tween-20 in Tris-buffered saline with 5% NFDM or 5% BSA). Primary antibodies were obtained from the following sources and used according to manufacturer’s recommendations; anti-b-actin (8457, 1:1,000, Cell Signaling Technology, Danvers, MA), anti-Chk1 (sc-8408, 1:250, Santa Cruz Biotechnology, Santa Cruz, CA), Anti-HA monoclonal antibody (HA.11 MMS-101R, 1:1000, Biolegand), Anti-H3.3 Total antibody (176840, 1:1000, Abcam), anti-H3.3 (31-1058-00, 1:1000, RevMAb Bioscience, South San Francisco, CA), Anti-Histone H3 mutated K27M antibody (ab190631, 1:500, Abcam), Anti-Histone H3.3 phospho S31 antibody (ab92628, 1:2000, Abcam), Anti-Tri-Methyl-Histone H3 Lys27 antibody (9733, 1:1000, Cell signaling) and anti-β-tubulin (ab21058, 1:5000, Abcam). The blots were then incubated with an anti-mouse or anti-rabbit IgG-HRP secondary antibody (7074, 7076 Cell Signaling), incubated with ECL solution (Amersham, GE Healthcare, Chicago, IL), and imaged on a AI600 Chemiluminescent Imager (Amersham). Densitometry was performed using ImageJ.

**Immunofluorescence.** Fixed cell immunofluorescence microscopy was performed as previously described[46]. Briefly, cells were fixed in methanol at −20 °C, blocked in TBS (50 mM Tris, pH 7.4,

150 mM NaCl), 3% bovine serum albumin (BSA) and 0.5% Tween-20 for 1 hr at RT. Primary antibody incubations were diluted in 3% BSA/TBS for 16 hr at 4°C. Primary antibodies and dilutions used are: anti-histone H3 K27M, (31-1175-00, 1:10,000, RevMAb Bioscience), anti-histone H3 K27me3 (ABE44, 1:10,000, Millipore), anti-histone H3.3 Ser31P (Ab62207, 1:5,000, AbCam, Cambridge, MA), anti-p53, (2524, 1:1,000, Cell Signaling, Danvers, MA), and anti-a-tubulin, (T9026, 1:1,000, Sigma-Aldrich, St. Louis, MO). Cells were washed three times for 10 minutes each and then indirect fluorescence was achieved using Alexa 488, 594, or 660 conjugated goat secondary antibodies (Life Technologies) for 3 hours at room temperature. Cells were washed three times for 10 minutes with 1% BSA/0.05% Tween TBS and incubated with 4′, 6-diamidino-2-phenylindole (DAPI) diluted in TBS for 5 min. Coverslips were mounted using ProLong Gold anti-fade (Life Technologies, Carlsbad, CA).

Fixed cell images were collected using a Leica DM RXA2 microscope (Leica Microsystems, Buffalo Grove, IL), equipped with a 63x 1.4NA apo objective and Hamamatsu ORCA–ER CCD camera (Hamamatsu, Bridgewater, NJ). Images were collected as a Z-series, with 0.25 μm spacing. All images were exported to Adobe Photoshop (Adobe, San Jose, CA) for final layout.

The measurement of H3.3 phospho-Ser31 fluorescence levels on isolated chromosomes was done as described[47], [48] modified here. Briefly, cells were immunolabelled with anti-H3.3 Ser31P. Coverslips were scanned to identify cells metaphase cells or anaphase cells with single misaligned chromosomes. A square region of interest (ROI) was drawn and replicated six times, each box placed non-overlapping over the pericentromere regions of metaphase cells or arms of mis-aligned anaphase chromosomes. Six ROI's of identical size were generated and placed in non-overlapping fashion away from the chromosomes; the fluorescence intensity in each region was then measured, the six regions averaged, and the non-chromosomal intensities subtracted from the chromosomal fluorescence.

**Live Cell Microscopy.** For live-cell imaging, cells cultured on biocleaned coverslips were assembled into custom-built imaging chambers [12], filled with Phenol Red-free DMEM supplemented with 12.5 mM Hepes buffer and 10% FBS.

Live-cell images were collected using a Yokogawa CSU-10 spinning disk confocal (Yokogawa Electric, Tokyo, Japan), with 200 mW 488nm diode laser (Coherent, Santa Clara, CA) and Hamamatsu 9100 EM-CCD camera mounted on a Leica DM RXA2 microscope equipped with a 63x 1.3NA glycerol immersion Apochromatic objective or a 20x 1.2 NA multi-immersion Apochromatic objective, housed in a custom-built Plexiglas box and maintained at 37°C with a proportional heat controller. All imaging systems were controlled by Simple PCI (Hamamatsu, Bridgewater, NJ) or Slidebook (3I, Denver CO) software. Time-lapse images were collected as a Z-series, with 2 μm spacing. For live cell chromosome mis-segregations Z-stacks were collected at intervals ranging from one to three minutes. To follow live cells through their full cell cycle, low magnification Z-stacks (20x magnification) were collected at intervals ranging from fifteen to thirty minutes. All images are presented as maximum projections. Cell cycle timing is calculated as the onset of anaphase in one mitotic division to the onset of anaphase in the next mitotic division.

**Growth Curves.** Cells were plated into 24 well dishes in DMEM with 10% FBS. Growth was monitored on an IncuCyte S3 in incubator microscope at 37°C and 5% $CO_2$. Cells were imaged in white light every 4 hours using a 10x lens. IncuCyte software was used to calculate percentage confluency and data exported in text form. Growth curves were then regraphed in Origin 2018b (OriginLab).

**Viral infections in vivo**: Nestin-TVA mice have been described [13]. Infected DF-1 cells from a confluent culture in a 10-cm dish were trypsinized, pelleted, resuspended in 50 μl HBSS (24020-117 Gibco), and placed on ice (8). Newborn N-TVA mice were injected intracranially 2 mm forward and to 2 mm right of the intersection of the coronal and sagittal sutures with 5μl of infected DF-1 cells using a gas-tight Hamilton syringe. Cohorts were injected with RCASBP(A) PDGFβ-P2A-H3.3$^{WT}$, PDGFβ-P2A-H3.3$^{S31A}$ or PDGFβ-P2A-H3.3$^{K27M}$. Censored survival data was analyzed using a log-rank test of the Kaplan–Meier estimate of survival. All mice were monitored for tumor development daily. All animal experiments were performed in compliance with 'Care and Use of Animals' in Association for Assessment and Accreditation of Laboratory Animal Care (AAALAC)-accredited facilities and approved by the University of Minnesota IACUC.

**Immunohistochemistry**: Brain tissue from injected mice was fixed in 10% formalin, embedded in paraffin and 5-μm sections were adhered to glass slides. The sections were stained with hematoxylin and eosin or left unstained for immunohistochemistry (IHC). Tissue sections were de-paraffinized and antigen retrieval was performed in 'Rodent Decloaking' buffer (Biocare Medical Pacheco, CA) by boiling for 10 min. Sections were treated with 3% $H_2O_2$ and blocked in Background Sniper (Biocare Medical) for 10 min. Primary antibodies were diluted in Renaissance background reducing diluent (Biocare Medical). Antibodies against the following antigens were used: GFAP (GA524, 1:5000, Dako); Nestin (ab6142, 1:200, Abcam); H3.3 K27M (ab6142, 1:500, Abcam); Ki67 (SP6 RM-9106, 1:200 ThermoScientific). Olig2 (AB9610, 1:2000, Millipore); H3 K27 me3 (9733, 1:200 Cell signaling). Sections were incubated overnight at 4°C and probed with Mach 4 rabbit polymer reagent (Biocare Medical). H3.3$^{WT}$, H3.3$^{K27M}$ and H3.3$^{S31A}$ expression was detected using an antibody to the HA epitope (HA.11 MMS-101R, 1:100, Biolegand) and the Mouse-on-Mouse HRP-Polymer kit and blocking regents (MM510L Biocare Medical). Visualization was carried out with DAB (Biocare Medical). Sections were counterstained with hematoxylin (IHC World, Ellicott City, MD).

**Statistics and reproducibility.** The numbers of cells analyzed are indicated in the legend to each figure. Graphs were generated in Origin 2018b. Statistical analysis was done using a two-tailed t-test in Microsoft Excel.
